# Improved Annotation with *de novo* Transcriptome Assembly in Four Social Amoeba Species

**DOI:** 10.1101/054536

**Authors:** Reema Singh, Hajara M. Lawal, Christina Schilde, Gernot Glöeckner, Geoff J. Barton, Pauline Schaap, Christian Cole

## Abstract

**Background:** Annotation of gene models and transcripts is a fundamental step in genome sequencing projects. Often this is performed with automated prediction pipelines, which can miss complex and atypical genes or transcripts. RNA-seq data can aid the annotation with empirical data. Here we present de novo transcriptome assemblies generated from RNA-seq data in four Dictyostelid species: *D. discoideum, P. pallidum, D. fasciculatum* and *D. lacteum*. The assemblies were incorporated with existing gene models to determine corrections and improvement on a whole-genome scale. This is the first time this has been performed in these eukaryotic species.

**Results:** An initial *de novo* transcriptome assembly was generated by Trinity for each species and then refined with Program to Assemble Spliced Alignments (PASA). The completeness and quality were assessed with the Core Eukaryotic Genes Mapping Approach (CEGMA) and Transrate tools at each stage of the assemblies. The final datasets of 11,315-12,849 transcripts contained 5,610-7,712 updates and corrections to >50% of existing gene models including changes to hundreds or thousands of protein products. Putative novel genes are also identified and alternative splice isoforms were observed for the first time in *P. pallidum, D. lacteum* and *D. fasciculatum*.

**Conclusions:** In taking a whole transcriptome approach to genome annotation with empirical data we have been able to enrich the annotations of four existing genome sequencing projects. In doing so we have identified updates to the majority of the gene annotations across all four species under study and found putative novel genes and transcripts which could be worthy for follow-up. The new transcriptome data we present here will be a valuable resource for genome curators in the Dictyostelia and we propose this effective methodology for use in other genome annotation projects.

## BACKGROUND

Whole genome sequencing projects are within the scope of single laboratories. The Genomes OnLine Database [1] reports (as of 13^th^ May 2016) there are 76,606 sequenced organisms, of which 12,582 are eukaryotes. However, only 8,047 are reported as being complete. Annotation of gene models is a requirement for a complete genome [2], There are several complementary strategies for achieving gene annotation in novel genomes including gene prediction[3, 4], expressed sequence tag (EST) libraries [5] and RNA sequencing (RNA-seq) data [6]. Gene prediction methods are limited in the complexity of the gene models they are able to produce; alternative splice sites are unpredictable and untranslated regions (UTRs) have subtle signals[7], EST libraries, if available, are usually fragmented and incomplete. RNA-seq data is dependent on good alignments to the reference. *De novo* transcriptome assembly is equally able to fulfil this function, although it can be computationally challenging [8] [9] [10]. Transcriptome assembly methods can be either reference-guided or reference-free [11] [12], Reference-guided methods have the advantage of simplifying the search space, but are dependent on the relevance, quality and completeness of the reference. Reference-free methods do not have any dependencies, but need to deal with sequencing errors sufficiently well to avoid poor assemblies [11–13]. We present the application of a *de novo* transcriptome assembly to four eukaryotic species: *Dictyostelium discoideum, Polysphondylium pallidum, Dictyostelium fasciculatum* and *Dictyostelium lacteum*. The genome of *D. discoideum* was published in 2005, it is 34 Mb in size and has been assembled into six chromosomes, a mitochondrial chromosome, an extra-chromosomal palindrome encoding ribosomal RNA (rRNA) and three ‘floating’ chromosomes [14], The genome was generated via dideoxy sequencing and contigs were ordered into chromosomes by HAPPY mapping (*D. discoideum*) [14] [15] [14] and still contains 226 assembly gaps.

In contrast, the similar sized genomes of *P. pallidum, D. lacteum* and *D. fasciculatum* were sequenced more recently using both dideoxy and Roche 454 sequencing. Their assembly was assisted by a detailed fosmid map and primer walking, leading to only 33 to 54 gaps per genome, but are more fragmented with 41, 54 and 25 supercontigs, respectively [15] (Gloeckner et al., 2016, submitted). The *D. discoideum* genome has been extensively annotated via the Dictybase project [16], whereas the gene models for P. *pallidum, D. fasciculatum* and *D. lacteum* available in Social Amoebas Comparative Genome Browser [17] are primarily based on computational predictions.

The social amoeba *D. discoideum* is a widely-used model organism for studying problems in cell-, developmental and evolutionary biology due to their genetic tractability allowing elucidation of the molecular mechanisms that underpin localized cell movement, vesicle trafficking and cytoskeletal remodeling as well as multicellular development and sociality. The social amoebas form a single clade within the Amoebozoa supergroup and are divided into four taxon groups according to molecular phylogeny based on SSU rRNA and α-tubulin sequences [15]. The four species under study here represent each of the four groups: *D. discoideum* (group 4), *P. pallidum* (group 2), *D. fasciculatum* (group 1) and *D. lacteum* (group 3). Genome annotations are not static and benefit from the application of additional evidence and new methodologies [7, 18]. Therefore we present, for the first time, substantially updated annotations based on a *de novo* transcriptome assembly for the *D. discoideum, P. pallidum, D. fasciculatum* and *D. lacteum* genomes.

## METHODS

### Sample preparation

Sequencing data were obtained from four RNA-seq experiments. The *D. discoideum* data were obtained from an experiment comparing gene expression changes between wild-type cells and a diguanylate cyclase (dgcA) null mutant at 22 h of development [19]. The *P. pallidum* data were obtained at 10 h of development in an experiment comparing wild-type and null mutants in the transcription factor cudA (Du, Q. and Schaap, P. unpublished results). In this experiment *P. pallidum* cells were grown in HL5 axenic medium (Formedium, UK), starved for 10 h on nonnutrient agar, and harvested for total RNA extraction using the Qiagen RNAeasy kit. The data for *D. lacteum* and *D. fasciculatum* were obtained from developmental time series [20]. For these series cells were grown in association with *Escherichia coli* 281 washed free from bacteria and plated on non-nutrient agar with 0.5 % charcoal to improve synchronous development. Total RNA was isolated using the Qiagen RNAeasy kit at the following stages: growth, mound, first fingers, early-mid culmination, fruiting bodies. *D. lacteum* RNAs were also sampled at three time points intermediate to these stages.

### Illumina Paired End Sequencing

Total RNA was enriched for messenger RNA (mRNA) using poly-T oligos attached to magnetic beads and converted to a sequencing ready library with the TruSeq mRNA kit (Illumina), according to manufacturer’s instructions and 100 basepairs (bp) paired-end sequenced using an Illumina HiSeq instrument. For the *D. discoideum* and *P. pallidum* samples, 1 μg of total RNA was used as starting material, with 4 ul of 1:100 dilution External RNA Controls Consortium (ERCC) ExFold RNA Spike-In Mixes (Life Technologies) added as internal controls for quantitation for the RNA-Seq experiment and sequenced at the Genomic Sequencing Unit, Dundee. In total there were 433M, 413M, 171M and 319M reads respectively for *D. discoideum, P. pallidum, D. fasciculatum* and *D. lacteum*.

### Data processing and *de novo* transcriptomics assembly

The quality of the raw reads was checked with FastQC [21] and the reads were found to have high quality scores across their full length. No trimming of the data was performed, as aggressive trimming can negatively impact on the quality of assemblies [22], All reads for each species were combined prior to *de novo* assembly. Being a more mature genome the *D. discoideum* data was used to verify the methodology, thereby giving a reference point for the other, less well characterised, species. Figure 1 shows a schematic of the overall workflow.

**Figure 1.**
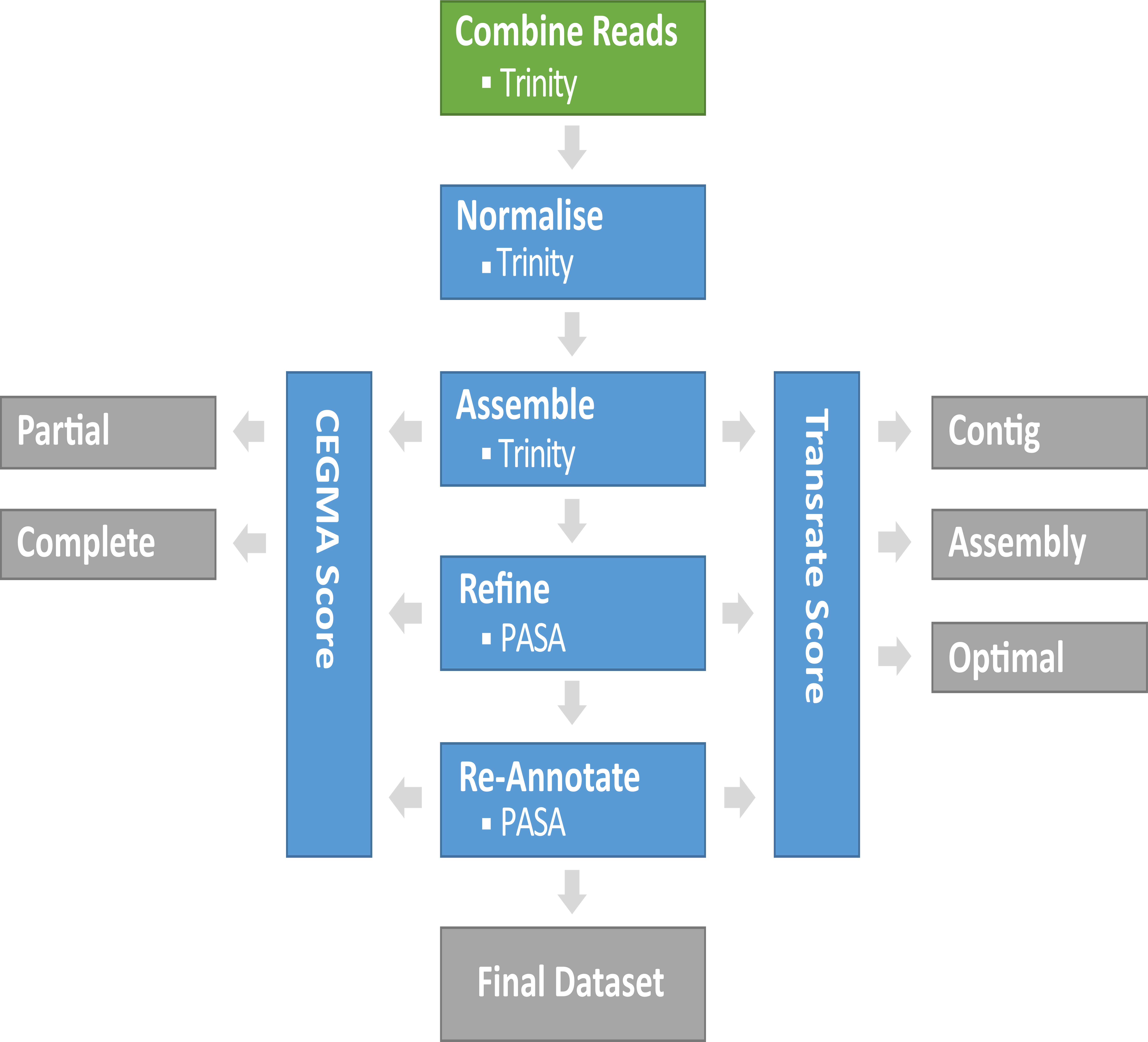
The *de novo* transcriptomics assembly workflow. The reads are input at the top in green, all computational steps are in blue and all data or quality control outputs are shown in grey. PASA is the Program to Assemble Splice Alignments tool[26]. See main text for description of PASA1 and PASA2 steps. CEGMA is the Core Eukaryotic Genes Mapping Approach tool[29].

Trinity version 2013.11.10 [8] was used for *de novo* assembly and normalisation of the read data was achieved with a kmer of 25 and aiming for 50x coverage of the genome. Following normalisation there remained 5.3M, 8.3M and 16.0M read pairs in *D. discoideum, P. pallidum* and *D. lacteum,* respectively. *D. fasciculatum* reads were not normalised as there were fewer than the recommended 300 million reads as per the Trinity manual. Trinity was run on the normalised reads using the - jacard-clip parameter and setting - k-min-cov to 4 in an attempt to reduce the number of fused transcripts in *P. pallidum* only. For the initial transcript set of *D. discoideum* and *P. pallidum* assemblies, any transcripts with BLAT (BLAST-like alignment tool) v35×l [23] hits to the ERCC spike-in sequences were removed from the *D. discoideum* and *P. pallidum* assemblies. *D. fasciculatum* and *D. lacteum* were not cultured axenically and thus the samples were contaminated by their bacterial food source. In order to remove the bacterial contamination *D. fasciculatum* and *D. lacteum*, transcripts were filtered with the TAGC (taxon-annotated GC-coverage) plot pipeline [24], TAGC determines for each contiguous sequence (contig) the proportion of GC bases, their read coverage and best phylogenetic match. With this information it is possible to identify which transcripts are mostly likely to be contaminants and removed. In order to remove the contamination, first all the transcripts were aligned to the BLAST ‘nt’ database using BLAST megablast. Using the trinity assembled transcripts, the BAM file of the reads mapped back to the transcripts and the transcripts to species mapping, non-target related transcripts were removed. The contaminant transcripts were differentiated on the coverage vs GC plots (see Supplementary Information and Figure S1).

The normalised set of reads were aligned with bowtie (0.11.3, with parameters applied as per Trinity script alignReads.pl)[25] to the whole transcript set and the total number of reads matching to each transcript were stored.

### Transcript Refinement

Program to Assemble Spliced Alignments (PASA) v2.0.0 [26] was used to refine the Trinity transcripts into more complete gene models including alternatively spliced isoforms. Initially developed for EST data, PASA has been updated to also work with *de novo* transcriptome data. Using the seqclean tool available with PASA, all the transcripts were screened and trimmed for low complexity regions, poly (A) tails and vector sequences. GMAP (Genome Mapping and Alignment Program) [27] and BLAT [23] were used to align the transcripts to their respective genomes. Trinity transcripts that failed to align to the already existing genome in both GMAP and BLAT were removed as ‘failed’. Remaining ‘good’ transcripts at this stage are termed the PASA1 dataset. Next, PASA takes existing annotations and compares them to the PASA1 dataset. PASA uses a rule-based approach for determining which transcripts are consistent or not with the existing annotation and updates the annotation as appropriate: new genes, new transcript isoforms or modified transcripts. PASA2 is the term used for the PASA assembled transcripts after updating with existing annotation.

### Assembly Quality Check

At each stage, the transcript datasets were assessed with Core Eukaryotic Genes Mapping Approach, CEGMA, (Parra, Bradnam et al. 2007) and Transrate v1.0.0 [13]. These methods take complementary approaches in assessing completeness and/or accuracy. Transrate v1.0.0 uses the read data and optionally the reference sequence as input. CEGMA defines a set of 248 core genes that are likely to be found in all eukaryotes. These genes are used as a proxy for minimum completeness based on the assumption that a eukaryotic genome or transcriptome assembly should encode a large proportion of the core set of genes. The CEGMA (Version 2.5) tool uses HMMs, defined for each of the core genes in the set, returning whether there are complete or partial matches within the *de novo* transcripts. Transrate calculates the completeness and accuracy by reporting contig score and assembly score. Contig score measures the quality of the individual contig, whereas assembly score measures the quality of whole assembly. CEGMA was run using default parameters for all the assemblies.

### Orphan RNAs

The full set of Trinity transcripts constitutes the best approximation of the assembly of transcripts expressed in the RNA-seq sequencing data. The transcripts were aligned against the existing genome and coding DNA (cDNA) references (from Dictybase (*D. discoideum*) and SACGB [17], (*D. fasciculatum, D. lacteum* and *P. pallidum*)) using BLAT. Any transcripts not matching the existing references were searched against the NCBI ‘nt’ database with BLAST [28] and with PSI-BLAST against the NCBI ‘nr’ database for the longest predicted ORF in any remaining transcripts without a match to ‘nr’. This exhaustive search allowed the categorisation of ‘annotated’ (transcript with match to known genome and/or cDNA), ‘known’ (match to related species), ‘artefact’ (match to non-related species (non-Dictyostelid)) and ‘putative novel’ (remainder) datasets.

### PCR and subcloning

*D. discoideum* genomic DNA (gDNA) was extracted using the GenElute mammalian genetic DNA extraction kit (Sigma). Polymerase chain reaction (PCR) reactions were run for 30 cycles with 50 ng of gDNA and 1 μM of primers with 45 s annealing at 55°C, 2 min extension at 70°C and 30 s denaturation at 94°C. The reaction mixtures were size-fractionated by electrophoresis, and prominent bands around the expected size were excised, purified using a DNA gel extraction kit (Qiagen) and subcloned into the PCR4-TOPO vector (Invitrogen). After transformation, DNA minipreps of clones with the expected insert size were sequenced from both ends.

## RESULTS AND DISCUSSION

### *De novo* Transcript Assembly

Table 1 shows a summary of the Trinity output for the *D. discoideum, P. pallidum, D. lacteum* and *D. fasciculatum de novo* transcriptome assemblies. Overall, the raw assemblies are similar in terms of total transcripts, GC content, and contig N50. *D. discoideum* is slightly anomalous in N50, mean length and transcripts ≥ 1,000 bp with all features being smaller than the other three assemblies. According to Dictybase the mean annotated gene length is 1,908 bp which is substantially larger than in the assembled transcripts (867 bp) suggesting that the *D. discoideum* transcripts are fragmented.

**Table 1.** Trinity assembly summary statistics

|  | *D. discoideum* | *P. pallidum* | *D. fasciculatum* * | *D. lacteum* * |
| --- | --- | --- | --- | --- |
| Total raw read pairs | 216,284,941 | 206,385,299 | 85,379,416 | 159,455,838 |
| Normalised read pairs | 5,278,467 | 8,273,023 | NA | 15,994,900 |
| *Assembled Statistics* |  |  |  |  |
| Total Trinity transcripts | 31,259 | 35,631 | 46,779 | 38,508 |
| Transcripts $\geq 1000$bp | 8,818 | 17,988 | 18,977 | 20,637 |
| GC content (%) | 28.2 | 35.7 | 35.6 | 31.2 |
| Maximum Transcript Length (bp) | 21,679 | 17,271 | 19,435 | 15,026 |
| *Stats based on all transcript contigs* |  |  |  |  |
| Contig N50 | 1,266 | 1,871 | 2,107 | 2,759 |
| Mean length (bp) | 871 | 1,338 | 1,212 | 1,671 |
| Total assembled bases | 27,302,238 | 47,819,726 | 56,707,337 | 64,358,120 |

\* Assembly statistics calculated after removal of bacterial transcripts (see Supplementary)

Figure 2 shows the distribution of transcript lengths for *D. discoideum, P. pallidum, D. fasciculatum, D. lacteum* (cyan) when compared to the available cDNA datasets (magenta). The *D. discoideum* cDNAs are manually curated, whereas the others are predicted. The transcript sets are enriched in short transcripts (<1000 bp) as compared to their cDNAs with the effect being most marked in *D. discoideum, D. fasciculatum* and *D. lacteum* (Figure 2A, Figure 2C and Figure 2D). The *P. pallidum* assembly is more similar to its cDNA reference dataset (Figure 2B). Interestingly, the longest assembled transcript in *D. discoideum* (21,679 bp) was found to be approximately half of the mitochondrial chromosome. We speculate that as the mitochondrion is gene rich and highly expressed that Trinity was unable to resolve overlapping reads from adjacent genes thereby joining them all into one ‘supercontig’.

**Figure 2.**
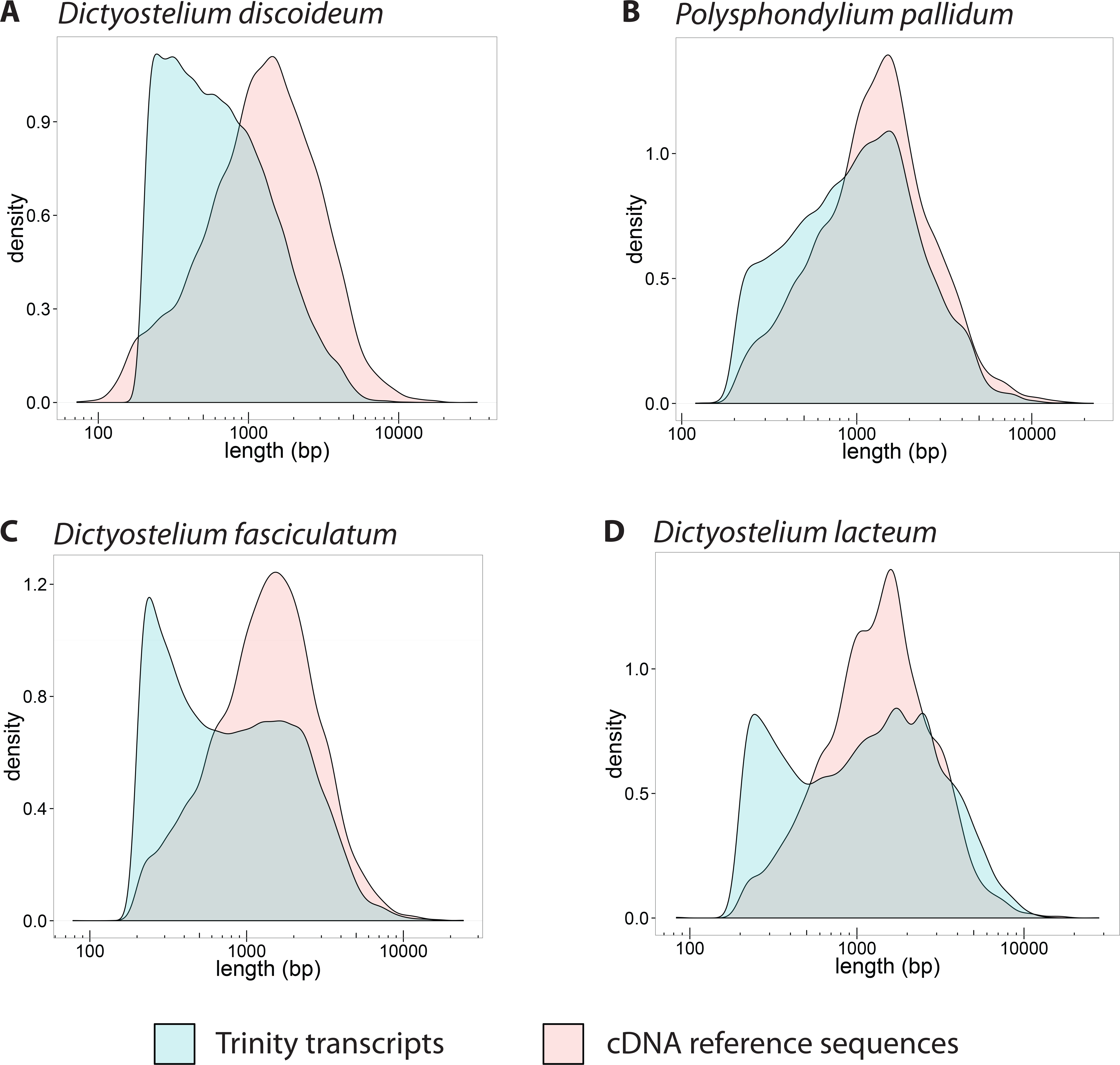
Trinity transcript length distributions. Comparison of assembled transcript sequence lengths (cyan) versus known cDN A sequence lengths (magenta) for *D. discoideum* (A), *P. pallidum* (B), *D. fasciculatum* (C) and *D. lacteum* (D).

The subsequent steps in the assembly were performed with PASA [26] which uses reference genome and transcript datasets to generate a refined and updated transcriptome assembly. The first stage (PASA1) takes the transcriptome assemblies, aligns them against the genome and clusters them into gene structures according to their genome alignments. Any transcripts, which do not align adequately to the genome are filtered out by PASA, under the assumption that they are misassemblies. In unfinished and complex genomes, it is possible there are missing gene loci in the genome reference. The missing loci may appear in a *de novo* transcriptome assembly and would be filtered out by PASA. The second stage (PASA2) uses the aggregated and filtered set of transcripts to refine the existing annotations for each of the species. At this stage, the gene models are updated with new or extended UTRs, new alternatively spliced isoforms are added and introns are added or removed. New genes are identified, and existing genes are split or merged as required by the *de novo* assembly data.

Table 2 shows the results of each stage of the assembly workflow from Trinity to each of the PASA steps and compared to the existing set of gene models from DictyBase (*D. discoideum*) or Augustus predictions (*P. pallidum, D. lacteum* and *D. fasciculatum*). It is clear that each stage the assemblies become more similar to the existing gene models (Table 2). For example, in all the species the total number of transcripts was 3-4-fold larger in the Trinity data than in the existing annotations. Although *de novo* assembly has the potential to identify novel genes and transcripts, a 3-fold increase is unlikely. By the end of PASA2, the transcript counts were within 1,500 of the existing models, with *D. fasciculatum, D. lacteum* and *P. pallidum* having more genes than in their Augustus-predicted models, and *D. discoideum* having 760 fewer genes than in the DictyBase-curated models. This is to be expected as the gene prediction algorithms are unlikely to have found all transcripts, whereas the *D. discoideum* curated set will include genes expressed under certain conditions only (e.g. developmental time points) that were not part of the experiment included here. Mean transcript lengths increased through the workflow. In particular, for *D. discoideum*, the mean Trinity transcript length was 871 bp and the final PASA2 length was 1,787 bp indicating that the high fragmentation observable by an excess of short transcripts (Figure 2) has been reduced. Similarly, the total number of identified exons was reduced from the initial Trinity dataset.

**Table 2.**
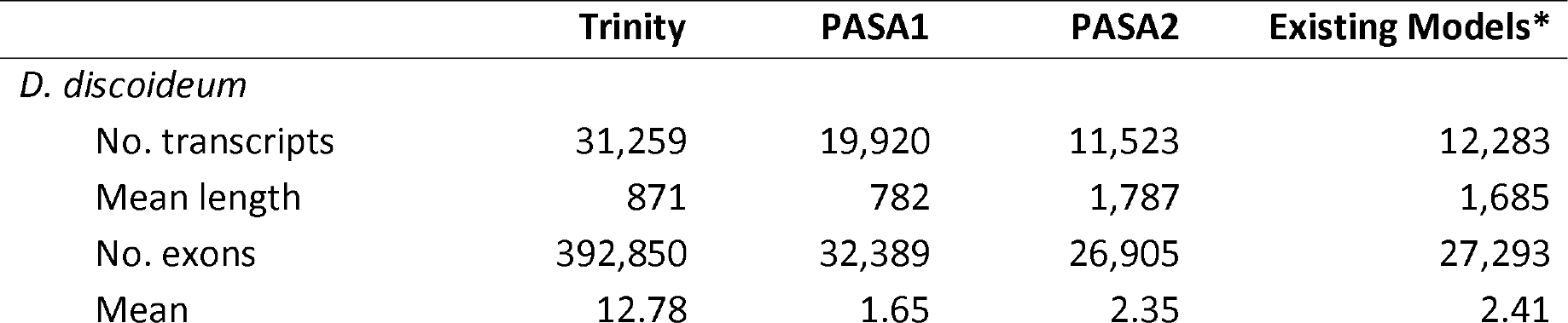
Comparison of transcript statistics at each stage of assembly.

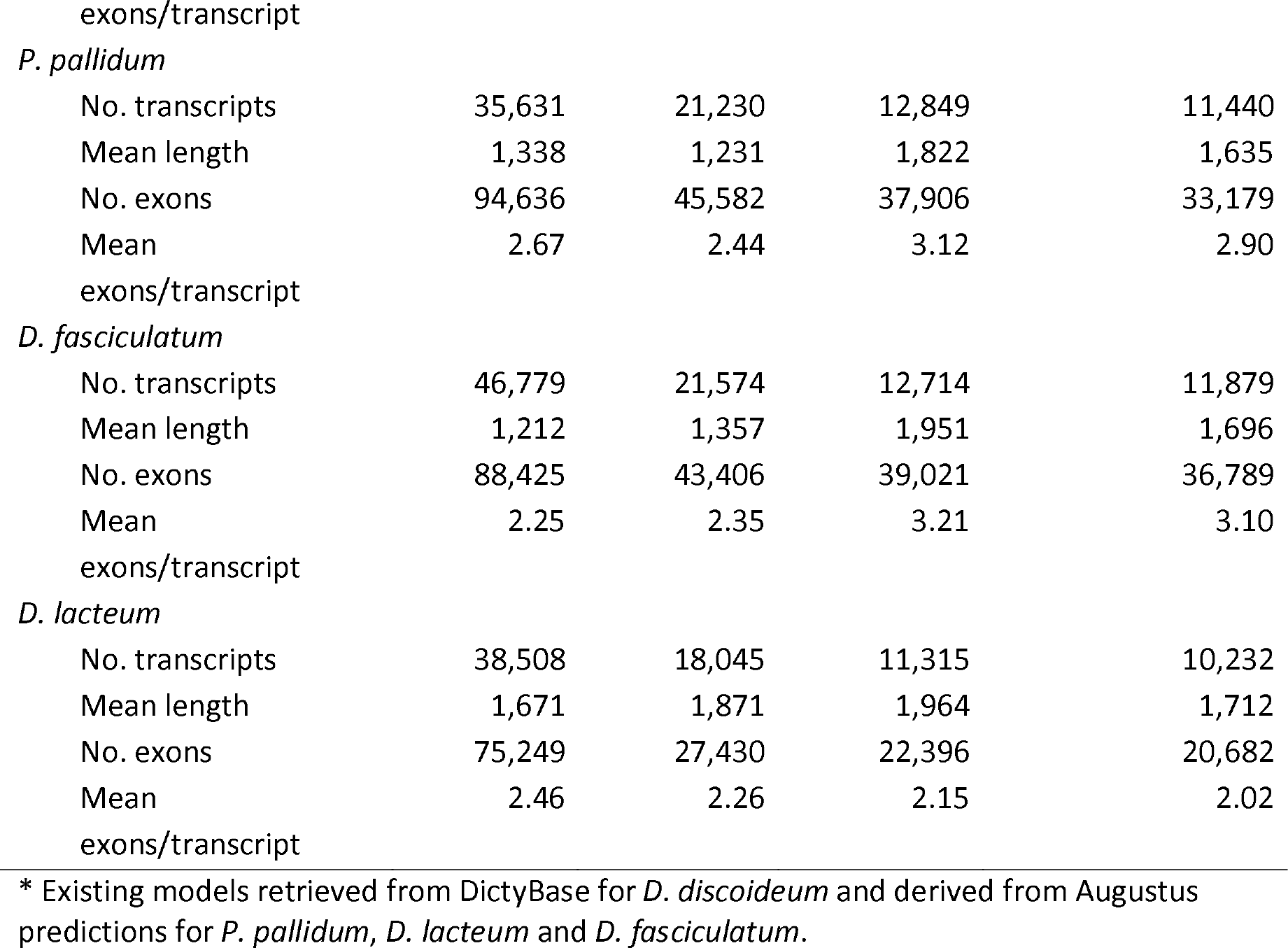

Overall, the initial Trinity assemblies have been refined from a fragmentary and redundant dataset to a more full-length and less redundant set of transcripts, which are more similar to the existing reference datasets in terms of total transcript counts, mean length, number of exons and exons per transcript (Table 2).

### Quality Assessment

Core Eukaryotic Genes Mapping Approach (CEGMA) and Transrate are tools which allow the assessment of completeness and accuracy of transcriptome assemblies. A set of 248 core eukaryotic genes (CEGs) was defined for CEGMA, for the purpose of assessing completeness in eukaryotic genomes [29]. CEGs are conserved across taxa and the majority should be present in the majority of eukaryotic species. A large fraction of missing CEGs could be indicative of an incomplete assembly. Figure 3 shows the comparison of complete and partial CEG matches in all four species for the genome reference, Trinity assembly, PASA1 refined transcripts and PASA2 updated annotations. In the ideal situation all CEGs would be detected in an assembly, however high sequence divergence or absence in the species of the CEGs will give lower maximum detection level. The whole genome CEGMA score represents the upper limit for any of the assemblies. All the datasets have >200 (>80%) complete or partial CEGs and are close the whole genome count suggesting the assemblies are nearly complete. It is noticeable, that the number of identified CEGs is consistently lower in the PASA1 data for all four species (Figure 3). This drop is due to the strict PASA filtering during transcript assembly. PASA1 only retains transcripts, which align to the reference with 95% identity and 90% length coverage. Manual checking of the CEGs that are identified in the Trinity data, but not in PASA1 reveals that they all are labelled as failed alignments. This suggests that either CEGMA is overly permissive in defining CEGs or that PASA1 is overly aggressive in filtering transcripts. PASA2 appears to ‘rescue’ this behaviour, presumably by including good annotations for genes that are poorly assembled in the Trinity data.

**Figure 3.**
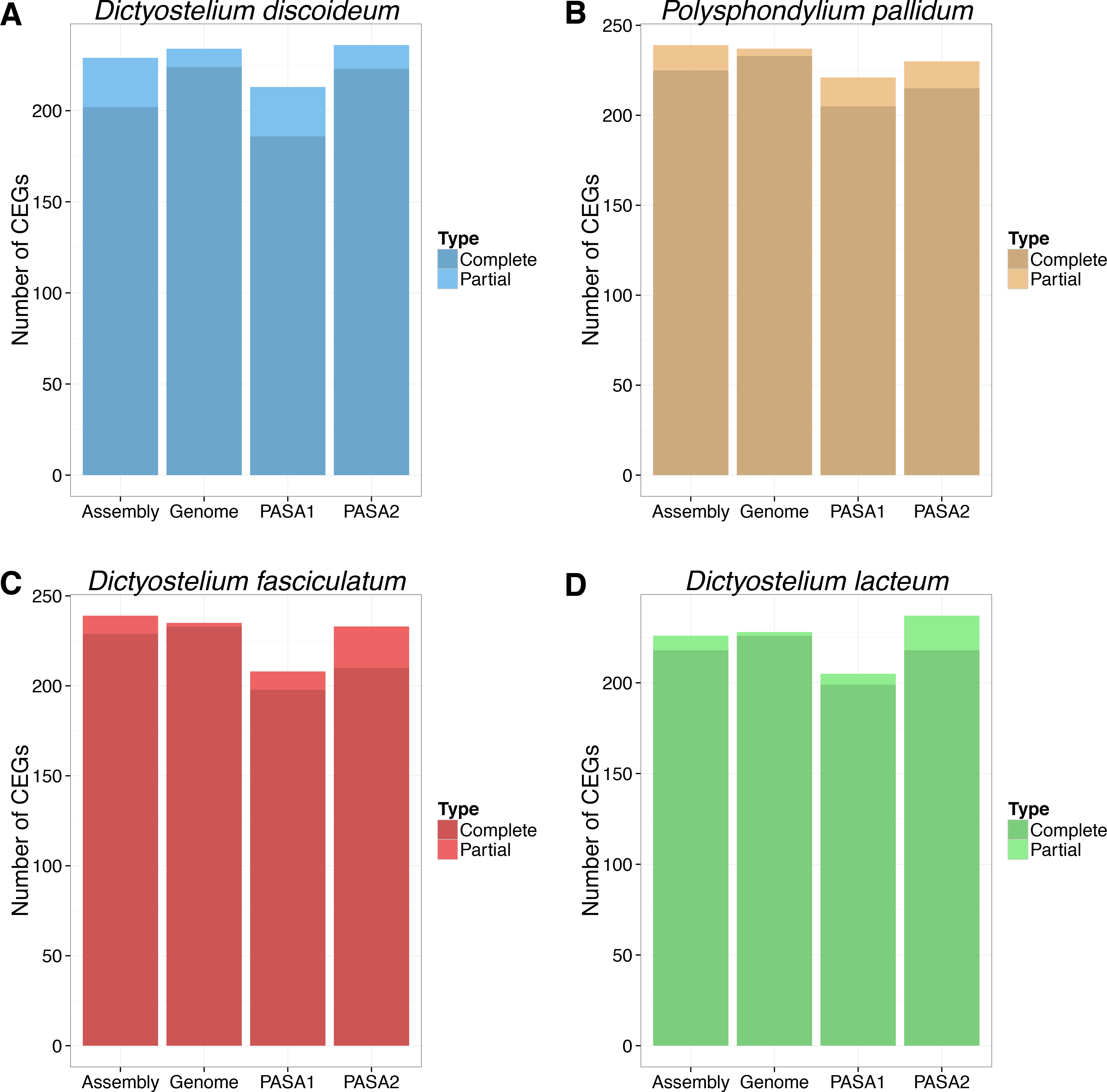
CEGMA complete and partial matches for *D. discoideum* (A, blue), *P. pallidum* (B< brown), *D. fasciculatum* (C, red) and *D. lacteum* (D, green) in the Trinity assembly, reference genome, PASA1 refined transcripts and PASA2 updated annotations.

Transrate assesses transcript quality by calculating several contig-level metrics based on the input RNA-seq data, and measures how well the read data support the contigs. Contigs are scored individually and then combined into an overall assembly score which ranges from 0 to 1. An optimal score is also reported, which predicts the best potential assembly score achievable by removing the worst scoring contigs in the dataset. An assembly score of 0.22 and optimised score of 0.35 was found to be better than 50% of 155 published *de novo* transcriptome assemblies [13]. A high Transrate score with a small improvement in the optimal score indicates a good *de novo* assembly, which is unlikely to be improved without further data or information.

Figure 4 compares the distribution of Transrate contig scores from the Trinity assembly, PASA1 refinement, PASA2 update and reference transcript/coding sequence (CDS) datasets for each of the four species. In contrast to the CEGMA data, the PASA1 data shows an improvement in Transrate contig scores when compared to the raw Trinity output meaning that the PASA1 transcripts are more consistent with the data, confirming that perhaps CEGMA is too permissive when assigning CEGs rather than PASA1 being too aggressive with its filtering. Notably the reference sequence datasets (‘CDS’ Figure 4) for *D. discoideum* and *P. pallidum*, show a lower median score than the PASA2 data, indicating that PASA2 is working well in combining the data with the existing annotations. There is little difference in *D. lacteum*. In *D. fasciculatum* the CDS data shows the best Transrate score of any of the assemblies.

**Figure 4.**
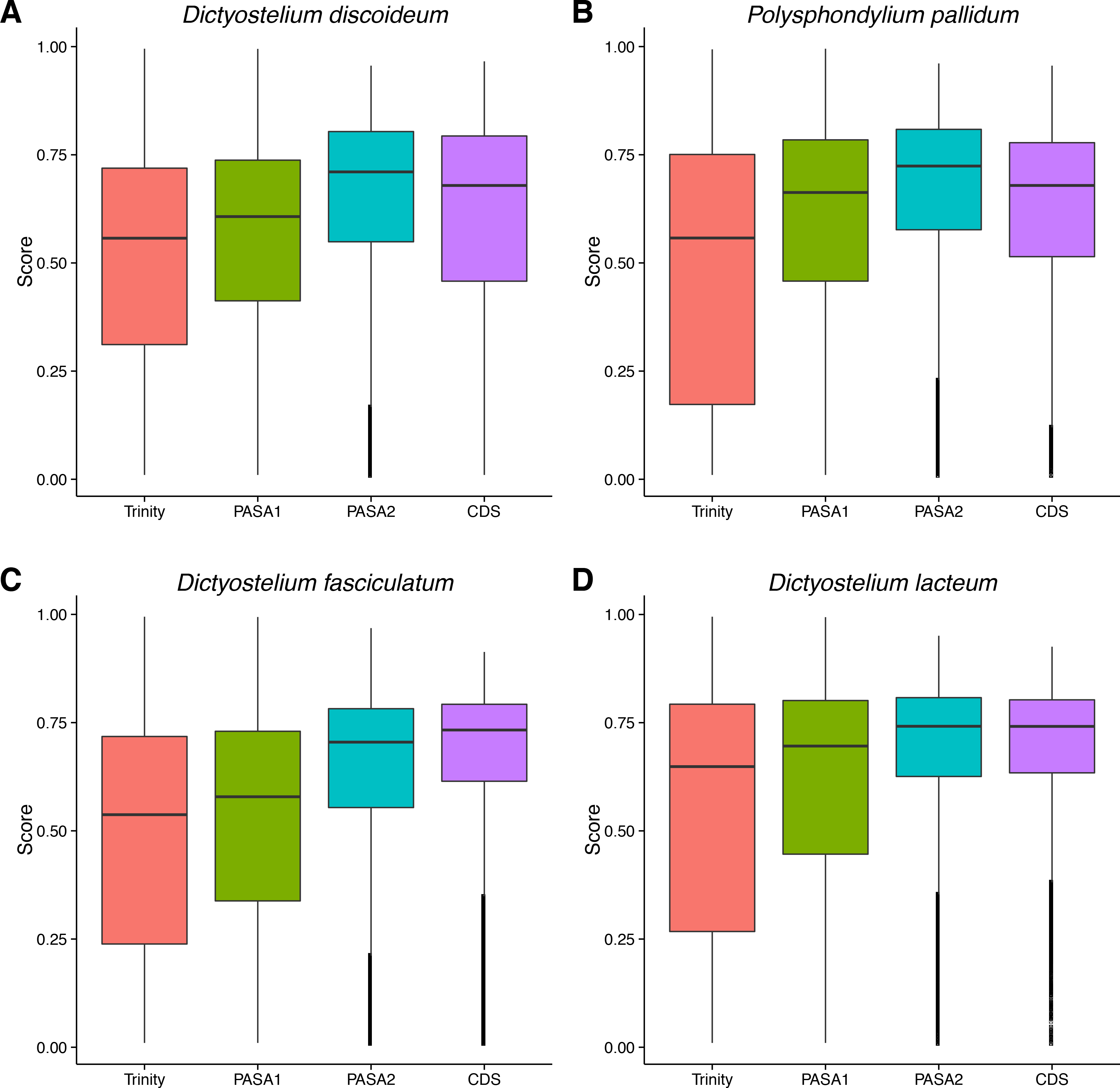
Distribution of Transrate contig scores (Score) [13] for the Trinity assembly [8], PASA1, PASA2 [26] and reference transcript (CDS) datasets for *D. discoideum* (A), *P. pallidum* (B), *D. lacteum* (C) and *D. fasciculatum* (D).

Figure 5 compares the Transrate assembly scores and optimal scores between PASA1 and PASA2 over the four species. The assembly scores range from 0.16 (*D. fasciculatum*) to 0.43 (*D. discoideum*) and the optimal scores range from 0.21 (*D. fasciculatum*) to 0.52 (*D. discoideum*). It is clear that PASA2 has much better Transrate scores (Figure 5A filled circles) than PASA1 (Figure 5A open circles), with all the PASA2 assemblies scoring better than 50% of published transcriptome assemblies (Figure 5A dotted black line). The optimal scores for PASA2 are also all better than 50% of published transcriptome assembly data (Figure 5A dotted cyan line), with the exception of *D. fasciculatum*. In *D. fasciculatum* the difference between the assembly (0.32) and optimal PASA2 scores (0.33) is small (Figure 5A green filled circles), suggesting that there is little improvement to the assembly possible given the read data for this species.

**Figure 5.**
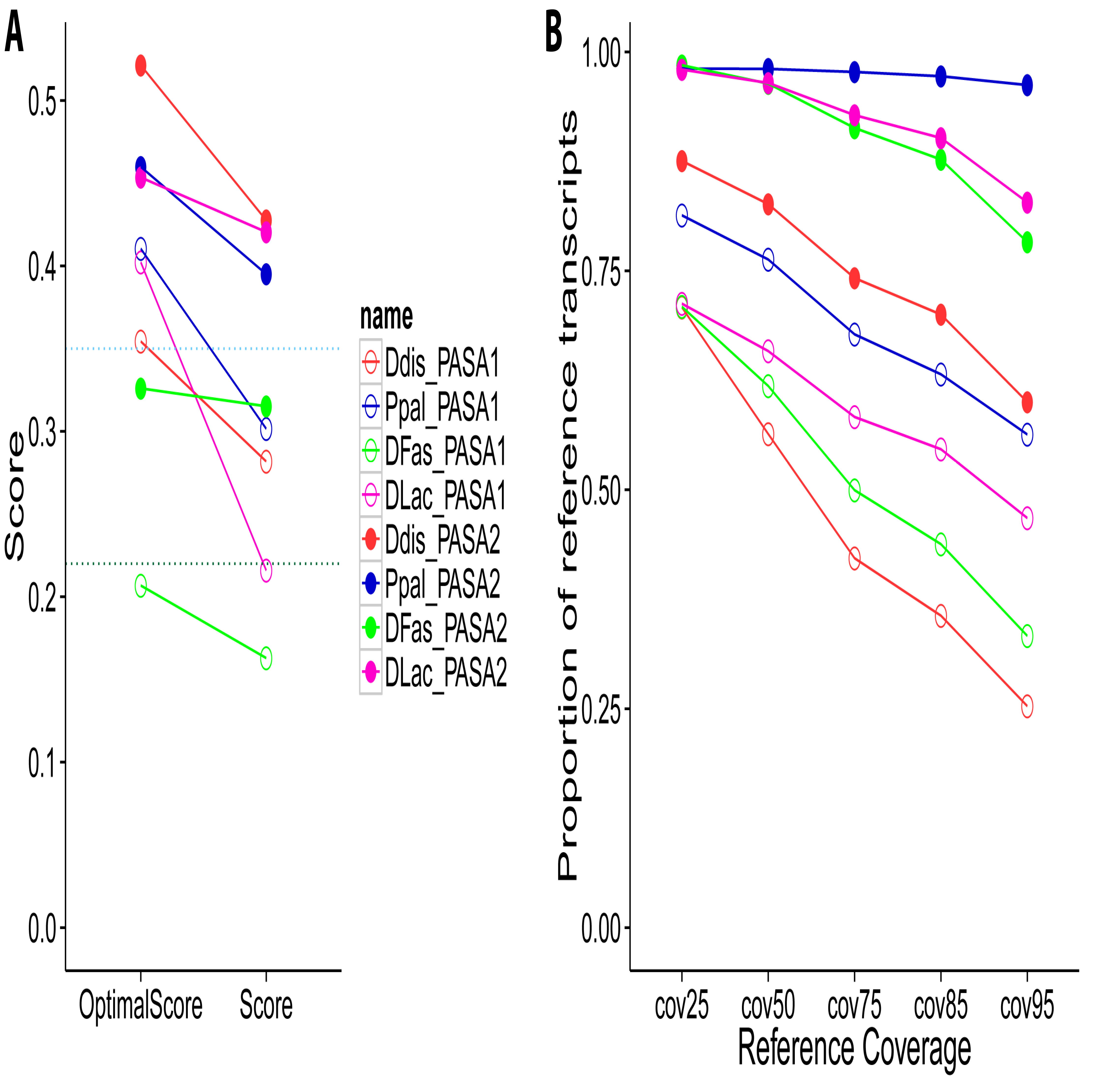
Transrate assembly scores and reference coverage metric. A) Compares the Transrate [13] assembly score and the optimised score between the PASA1 and PASA2 [26] steps in the four species (Ddis: *D. discoideum*, Ppal: *P. pallidum*, DFas: *D. fasciculatum*, DLac: *D. lacteum*). The dotted lines represent the Transrate scores that would be better than 50% of 155 published *de novo* transcriptomes as found by Smith-Unna and co-workers [13]: 0.22 overall score (black horizontal dotted line) and 0.35 optimal score (cyan horizontal dotted line). B) The proportion of reference protein sequences covered by assembled transcripts following PASA1 and PASA2 steps by at least 25%, 50%, 75%, 85% and 95% of the reference sequence length.

Transrate additionally has a reference-based measure, which aligns the transcripts to the reference protein sequences and the results are shown in Figure 5B. The y-axis in Figure 5B shows the proportion of reference sequences covered with transcript sequences at several thresholds (25%, 50%, 75%, 85%, 95%) of the reference. As before, Figure 5B highlights the improvements in PASA2 (filled circles) over PASA1 (open circles) with all species showing a higher proportion of reference sequences represented in the transcript data. The *P. pallidum* PASA2 assembly recapitulates the existing reference almost completely to 95% percent coverage (Figure 5 blue filled circles).

### Interpretation

What does an RNA-seq-based *de novo* assembly achieve when there is an already existing annotation either manually curated or generated via prediction? Is it worth it?

Table 3 details the results following PASA refinement of the existing gene models. Despite being a manually curated genome, the *D. discoideum* gene models where extensively modified by PASA with 7,182 being updated. Most of the updates in *D. discoideum* (6,750, 94%) are the result of UTR additions at 5’ and 3’ ends of genes, which were mostly missing in the existing models. The assemblies in the other species have a similar number of updates, but UTR-only updates to transcripts are a smaller fraction of the total. 187 new alternatively spliced transcripts, in 170 genes, were identified in *D. discoideum* (Table 3). There are currently 70 alternatively spliced transcripts, in 34 genes, annotated in Dictybase so this new data represents a 2.7-fold increase in the number alternatively splice transcripts and a 5-fold increase in genes. This number in *D. discoideum* could be an underestimate as the *D. fasciculatum, D. lacteum* and *P. pallidum* assemblies all have ~1000 alternate splice isoforms.

**Table 3.** Summary data following PASA transcript refinement and re-annotation.

|  | *D. discoideum* | *P. pallidum* | *D. fasciculatum* | *D. lacteum* |
| --- | --- | --- | --- | --- |
| Gene model updated | 7,182 | 7,712 | 6,664 | 5,610 |
| New alternate splice isoforms | 187 | 1,321 | 842 | 1,088 |
| Novel genes | 44 | 175 | 19 | 21 |
| Update results in modified protein | 554 | 2,252 | 5,393 | 4,741 |

Figure 6 provides examples, in each of the four species, of changes to the transcript models determined by PASA that are well supported by all the data. Each panel highlights a different type of change to the reference model. Gene DDB_G0295823 has a single transcript (DDB0266642 Figure 6A) with two exons and a single intron. The RNA-seq data (brown), Trinity assembly (purple) and PASA1 refinement (red) identifies extensions to the model, adding 5’ and 3’ UTRs to the annotation (green, narrow bars). The Trinity transcript (purple) is on the opposite strand to the reference transcript (black) and is corrected by PASA (red & green). The example in *P. pallidum* (Figure 6B) shows three new alternatively spliced products of the gene (Figure 6B green bars 1, 2, 3 labels). The three new models have the same coding region, but differ in their 5’-UTRs: two with differently sized introns and one without an intron. The new models also include a longer second coding exon (Figure 6B arrow), which increased the sequence of the protein product by 9 amino acids. Figure 6C shows an example, in *D. fasciculatum*, where a alternatively spliced transcript alters the protein product. The alternatively spliced isoform (Figure 6C, labelled 1) removes the first intron and extends the 5’-UTR when compared to the updated gene model (labelled 2). The CDS is shortened by 45 amino acids with the use of alternate start site, but the rest of the protein is identical. In the RNA-seq data it appears that this new alternative transcript is not the dominantly expressed isoform in the context of the whole organism. The final example is the merging of two *D. lacteum* genes into one (Figure 6D). The black bars show two distinct genes (DLA_11596 and DLA_04629), but the RNA-seq data (brown) and the Trinity assembly (purple bars) show uninterrupted expression across the intergenic region between the two genes (arrow). The PASA refinement and re-annotation (red and green bars) encapsulate the expression as a contiguous region with the coding region being in-frame over the two existing gene models. The annotation for the upstream DLA_11596 gene in SACGB [17] gives its best bi-directional hit in Uniprot/TrEMBL as *gxcN* in *D. discoideum* (DDB0232429, Q550V3_DICDI). *gxcN* codes for a 1,094 amino acid protein where DLA_11596 codes for a 762 amino acid protein and their pairwise alignment of DLA_11596 with DDB0232429 shows no overlap over the C-terminal 300 residues. The PASA2 gene fusion of DLA_11596/DLA_04629 (Figure 6D) codes for a longer, 1,029 protein which aligns across the full length of DDB0232429 in a pairwise alignment. We suggest that the existing gene model, DLA_11596, is a truncated form of a *D. discoideum gxcN* orthologue and that the fusion with the the downstream DLA_04629 gene represent the more accurate gene model.

**Figure 6.**
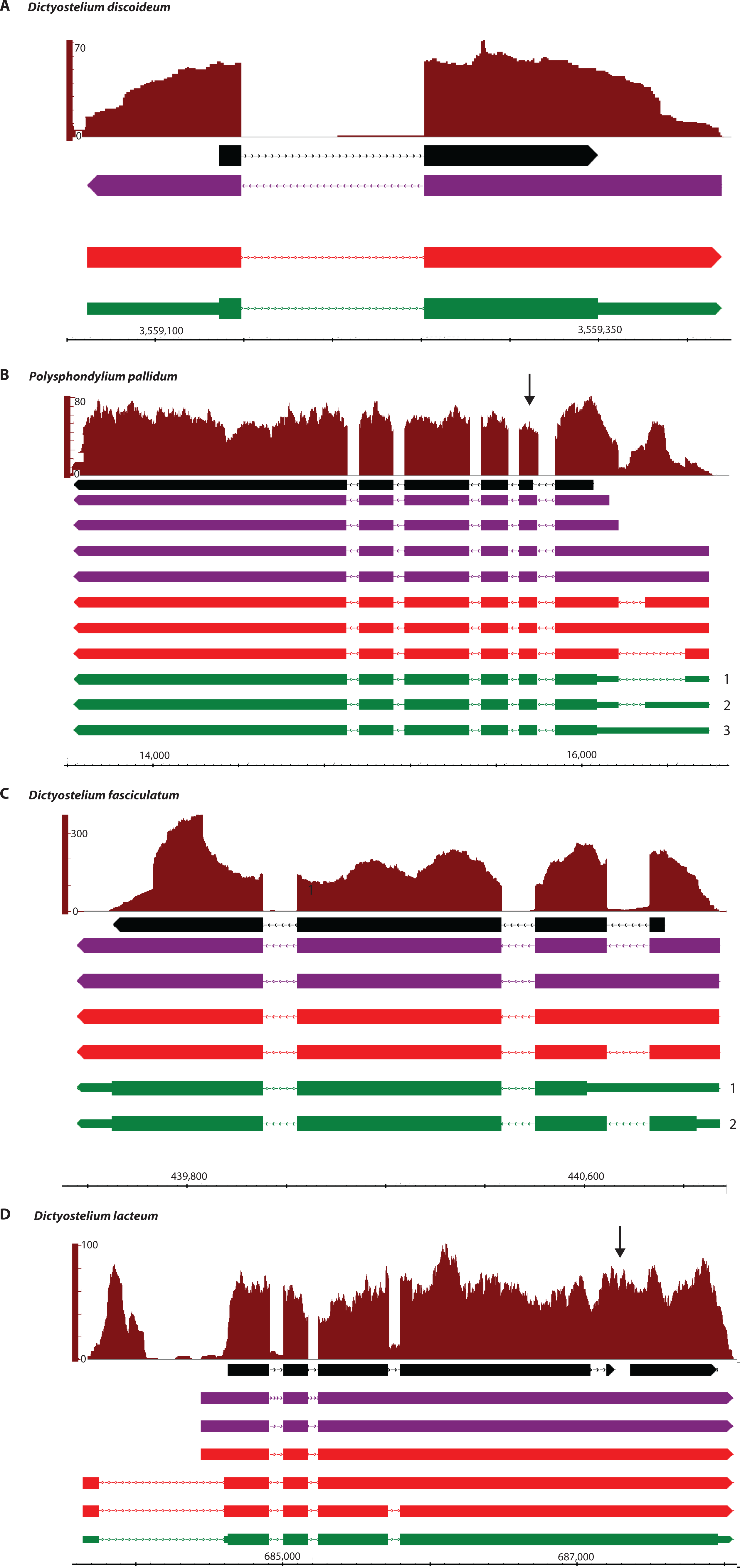
Examples of updated annotation in each species. Panels A-D compare the existing gene model (black bars) to pile-up of aligned RNA-seq reads (brown), Trinity de *novo* transcripts (purple bars), PASA1 refinement (red bars), PASA2 update (green bars). Intronic regions are shown by lines and UTRs by thinner green bars. The DNA strand is depicted by triangles at the end the bars: left end for reverse strand, right end for forward strand. Genes shown are: A) DDB_G029582 (*D. discoideum*), B) PPL_00079 (*P. pallidum*), C) DFA_02662 (*D. fasciculatum*) and D) DLA_11596/DLA_04629 (*D. lacteum*).

Given that *D. discoideum* has been extensively studied and the annotation curated by Dictybase, it is of note that our pipeline identified putative changes which altered the protein sequence of 554 genes (4.5% of total reference models) (Table 3). *D. discoideum* has been the focus of many functional studies including about 400 deletions in genes that are required for normal multicellular development (Gloeckner *et al,* under revision) [30]. Comparing the 554 *D. discoideum* genes with modified proteins to the developmentally essential genes, we found 16 genes (2.9%) that overlapped (see Supplementary Figure S3 for domain diagrams). Out of the 16, nine are either truncated or extended at the N-or C-terminal with probably little effect on protein function. In the remaining seven proteins, there is loss or gain of exons. Five proteins were updated with additional exons: DDB_G0268920, DDB_G0269160, DDB_G0274577, DDB_G0275445 and DDB_G0277719, and two proteins have an exon deletion: DDB_G0271502 and DDB_G0278639.

Investigating these protein changes in more detail revealed some errors in the underlying genome sequence, which resulted in some unusual gene models. Figure 7 shows clcD (chloride channel protein, DDB_G0278639) as an example. In the domain architecture of clcD, there are two CBS (cystathionine beta-synthase) domains present at positions 827-876 and 929-977 in the transcript sequence. In the updated sequence the protein is truncated and these two domains have been removed. This is likely to be incorrect since as all eukaryotic CLC proteins require the two C-terminal CBS domains to be functional [31]. How did this change occur in the *de novo* transcript assembly? In the existing annotation, there is an impossibly short two-base intron between the CLC domain and first CBS domain. Splicing requires a two-base donor and a two-base acceptor at either end of the splice site meaning at least four bases are required, not including any insert sequence. Investigating the RNA-seq genome aligned reads carefully reveals a single-base insertion immediately after the intron in 22/23 reads overlapping the region (yellow inset, Figure 7). The RNA-seq data turns the two-base intron into a three-base, in-frame codon inserting an isoleucine into the protein sequence and retains the CBS domains. By implication there is a missing base in the genome reference, which interrupts the open reading frame with a premature stop upstream of the CBS domains (arrow, Figure 7). PASA cannot deal with missing bases in the reference and erroneously truncates the, now out-of-frame, coding region four codons downstream of the missing base at a TGA stop codon. It also cannot create an impossible intron, which a human annotator presumably added in order to keep the transcript in-frame and retain the conserved CBS domains. PASA did make an error updating this gene, but it does not seem possible for it to have dealt with the missing base any other way.

**Figure 7.**
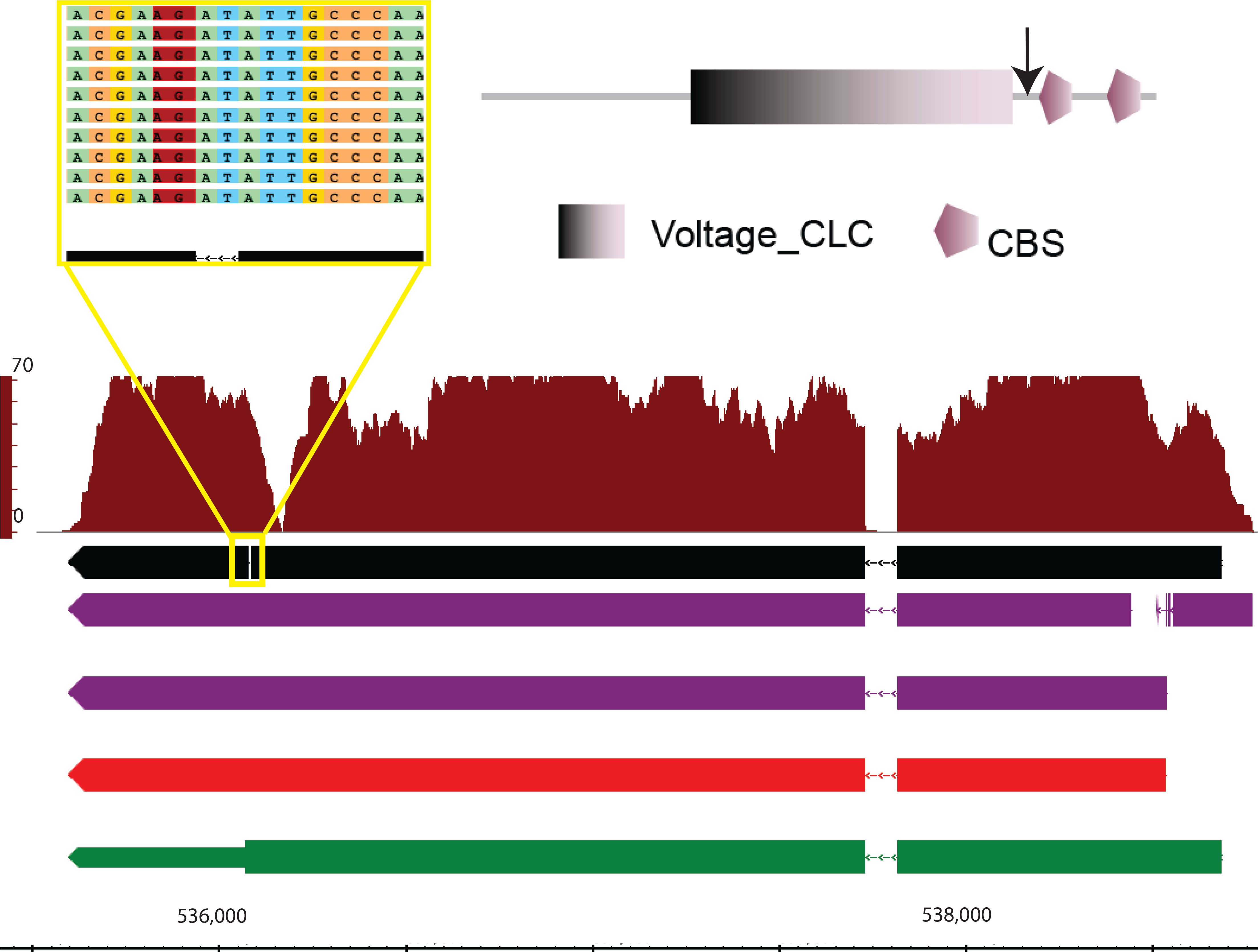
PASA update of the *clcD* locus (DDB_G0278639). See **Error! Reference source not found.** for meaning of coloured bars. Boxed in yellow, zoom in of RNA-seq reads covering Dictybase annotation of two-base intron. Reads are coloured by base, except in red highlights a region with an inserted base. Top right, SMART [43] protein domain architecture. Arrow shows the protein position of the yellow boxed region.

Inspection of all the *D. discoideum* gene models identified 119 sites in 102 genes with introns shorter than 5bp (see Supplementary Table S1). Of these genes, five have three tiny introns each. Four of them are either in poorly expressed genes or in poorly expressed regions within genes. One gene (DDB_G0279477), however, is well expressed across the full length. The gene contains two 3 bp introns and one 1 bp intron. The two 3 bp introns contain a TAA sequence encoding a stop codon, but according to the RNA-seq data the codons should be TTA (Leu) with evidence from 56 and 33 reads in the two sites, respectively. The 1 bp intron region is covered by 38 reads and one would not expect to see introns in RNA-seq data, by definition, it does seem highly unlikely for a 1 bp intron to exist given our current knowledge of mRNA splicing: canonical GU-AG dinucleotides and a branch point >18 bp upstream from the 3’ splice site. For this gene, there are clear errors in the genome sequence, which have lead to the creation of an erroneous gene model to compensate for them. It is arguable that none of the 119 <5bp introns are genuine but are artificial constructs to fix problems with the gene models. We recommend that gene annotators revisit these genes and consider updating the models [32] [7] and the underlying genome using RNA-seq data as evidence[33, 34].

The protein changes in *D. fasciculatum, D. lacteum* and *P. pallidum* number in the thousands (Table 3) highlighting that computational gene prediction is only a first step in annotating a genome. A reliable genome annotation requires evidence from many sources of information [18]. The types of protein changes seen in these three species range from inappropriately fused or split genes (see Figure 6 bottom panel for an example) via insertions/deletions to changes in protein coding start/stop codons positions resulting in extended or truncated coding sequences. All the PASA2 outputs are in the form of GFF files viewable within any genome browser. We have made an IGB Quickload server available for easy browsing of the data (http://www.compbio.dundee.ac.uk/Quickload/Dictyostelid_assemblies).

In the *D. discoideum, D. fasciculatum, D. lacteum* and *P. pallidum* datasets 44, 19, 21 and 175 novel putative genes were identified by PASA respectively (Table 3). These novel genes are in genomic loci with no current annotated gene model or where an existing model is substantially modified. The 44 *D. discoideum* novel genes, defined by 47 transcripts, were examined by eye in IGB [35] against all known *D. discoideum* reference datasets, including predicted gene models (see Table S2). Of the 47 transcripts, 8 are novel alternate splice transcripts (Table S2). Although ‘novel’ suggests there is no existing annotation at the locus of interest, if a gene update is sufficiently different from the reference gene model, PASA may consider that locus as a novel gene. In most of these cases the new transcript represents a corrected model for a previously computationally predicted gene. Many of the predicted gene models were annotated in Dictybase as as pseudogenes and were originally ignored by PASA2, which only considers protein coding genes. Fragments of the pseudogenes do encode ORFs and PASA has reported them as being novel genes (Table S2), but it is not possible to be sure whether the protein products are expressed *in vivo* with this data. Out of the 47, it appears only 6 are truly novel as they do not overlap any previously annotated transcripts: novel_model_13, novel_model_23, novel_model_30, novel_model_31, novel_model_38 and novel_model_39. All bar model_model_23 have a sequence match to existing genes, suggesting that they are paralogues. The longest novel unannotated model is 510 AA in length (novel_model_31) and appears to be a duplicate copy of the leucine rich repeat protein IrrA present on the chromosome 2.

Notwithstanding the large number of updates to the existing *D. discoideum* annotations it is clear from Table 3 that there are substantially more changes in the other three species. In particular, the numbers of modified protein sequences are 4, 9 and 10-fold larger in *P. pallidum* (2,252), *D. lacteum* (4,741) and *D. fasciculatum* (5,393), respectively. Similarly, there are 7, 5 and 6-fold more novel alternate splice isoforms in the three species, respectively. For *P. pallidum* (1,321), *D. lacteum* (1,088) and *D. fasciculatum* (842), the gene models were predicted with Augustus (G. Glöckner, *personal communication*) which, given the updates found with PASA, suggests that although the predicted gene models are in the correct locus, many are inconsistent with empirical RNA-seq evidence. With respect to novel genes annotated by PASA, it is notable that *D. fasciculatum* and *D. lacteum* have fewer than either *D. discoideum* or *P. pallidum.* It is unclear why this would be. Many genes were inspected by eye with IGB [35] and overall the annotations appear appropriate, but there are many occasions where human intervention would make further improvements.

### Orphan RNAs

As mentioned above PASA requires that transcripts align to the genome before it can consider them for further analysis. It makes sense to use the genome as a filter for valid transcripts, however this makes the assumption that the genome is complete. Any gaps in the genome that include genes will result in filtering out perfectly valid transcripts.

To determine whether this has happened here, we isolated the transcripts that did not align to the genome and used a process of elimination to identify those transcripts that could be genuine. Table 4 breaks down the number of orphan RNAs and whether they match non-dictyostelid genes (‘artefact’), genes in other dictyostelid (‘known’) or neither (‘novel’). *D. fasciculatum* and *D. lacteum* have far more non-genome transcripts (6,559 and 6,465, respectively) than *D. discoideum* (69) or *P. pallidum* (26). This is likely due to the fact that these species, which were cultured on bacteria, contain chimeric misassemblies of bacterial and dictyostelid transcripts. Despite this, they still have 525 and 945 ‘known’ transcripts which have sequence matches to other Dictyostelids, higher than seen in *D. discoideum* (14) and *P. pallidum* (82). These transcripts are probably the best candidates for experimental assessment as genuinely non-genome transcripts.

**Table 4.** Annotation of Trinity transcripts.

|  | *D. discoideum* | *P. pallidum* | *D. fasciculatum* | *D. lacteum* |
| --- | --- | --- | --- | --- |
| Annotated | 28,698 | 29,405 | 32,875 | 28,463 |
| *Aligned to reference* |  |  |  |  |
| Known | 672 | 170 | 3,028 | 240 |
| Novel | 1,277 | 4,505 | 2,978 | 2,040 |
| Artefact | 98 | 376 | 288 | 134 |
| *Not aligned to reference* |  |  |  |  |
| Known | 14 | 82 | 525 | 945 |
| Novel | 69 | 26 | 6,559 | 6,465 |
| Artefact | 431 | 81 | 208 | 366 |

We further investigated the 69 *D. discoideum* ‘novel’ with a more sensitive PSI-BLAST search on their longest ORFs and queried their cognate proteins for functional domains using SMART. Table 5 shows the 11 most interesting hits based on the sequence match, read count and ORF length. They are all well expressed and have ORF lengths consistent with functional proteins. Three novel transcripts (comp4660_c0_seql, comp4660_c4_seql and comp5569_c2_seql) show similar sequence matches to DDB_G0292950 via PSI-BLAST searching, in spite of very low sequence similarity between them. DDB_G0292950 codes for a hypothetical protein which is not conserved in other dictyostelids and is poorly expressed (RPKM <1) at all time points in dictyExpress [36]. The three transcripts match across different parts of DDB_G0292950 indicating that they are different parts of the same larger gene. All transcripts identified in Table 5 were selected for experimental validation via PCR amplification, 8/11 were confirmed.

**Table 5.**
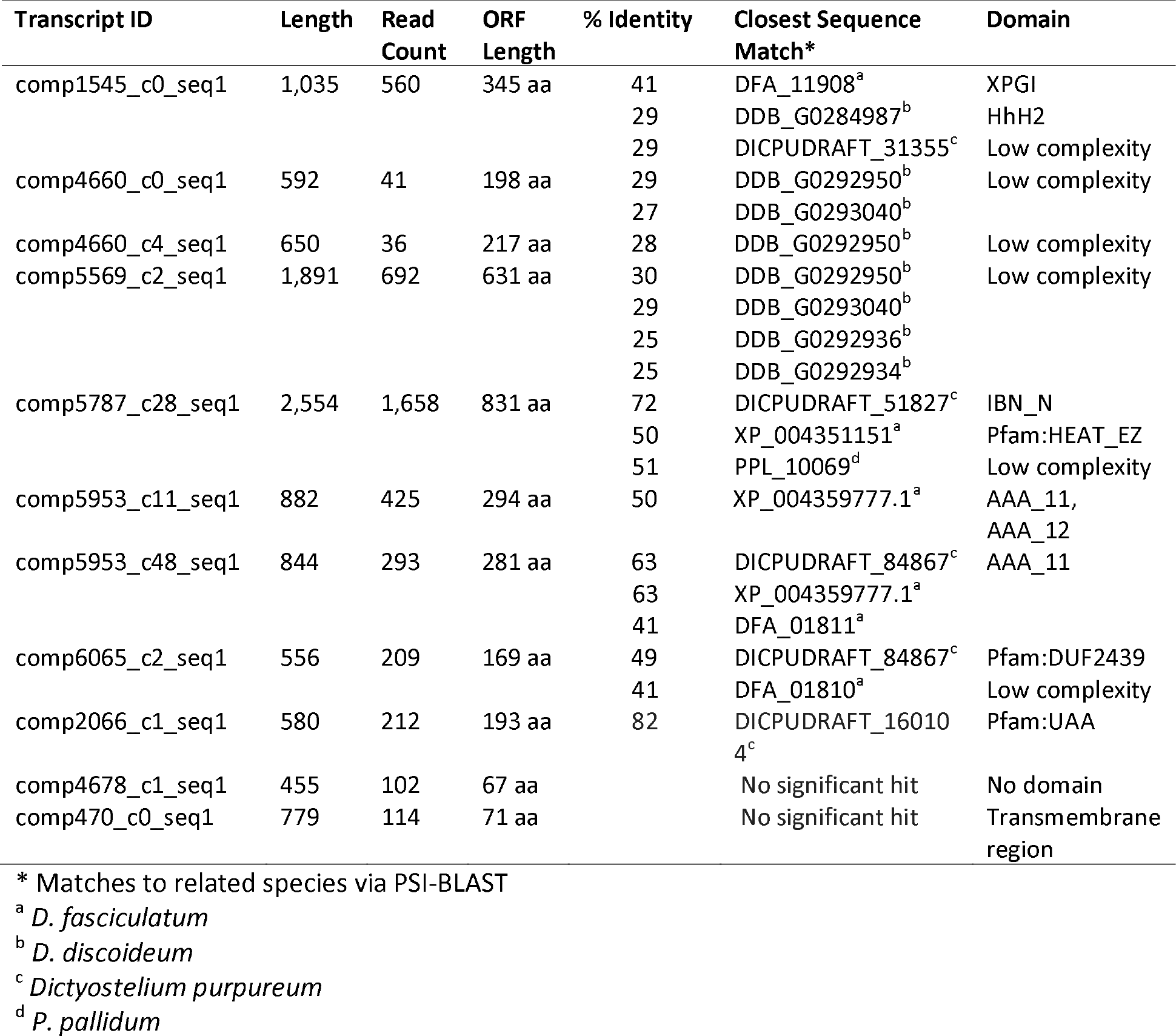
Homology and functional information for novel transcripts in D. discoideum.

The comp5787_c28_seql transcript has putative homologues in *D. fasciculatum, D. purpureum* and *P. pallidum* as shown in Figure 8. Sequence conservation is high as well as conservation of the Importin-beta N-terminal domain (IBN_N) and HEAT-like repeat (HEAT_EZ) domain architecture although the *D. discoideum* sequences appears to have an additional HEAT repeat domain (Figure 8).

**Figure 8.**
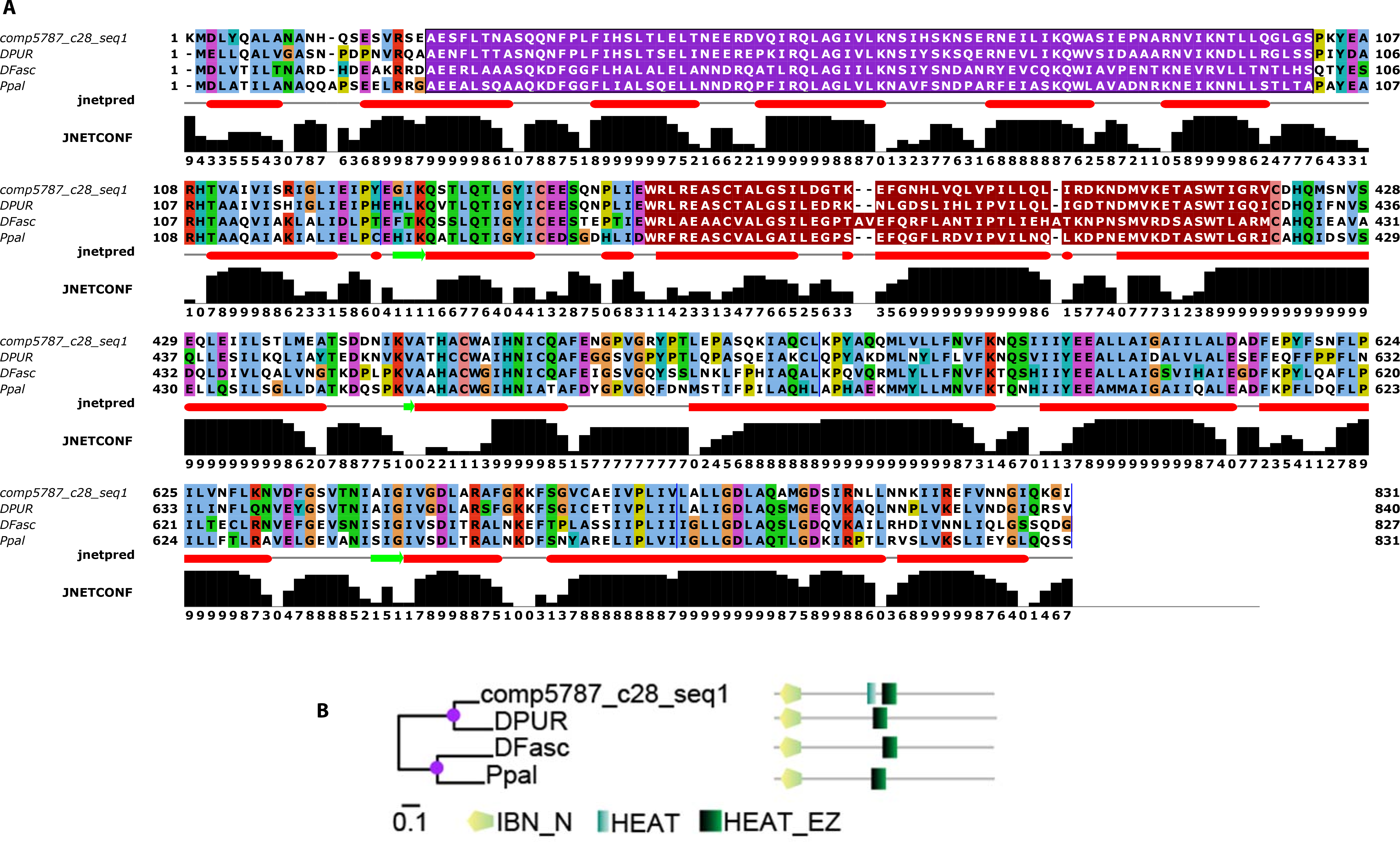
Protein sequence for comp5787_c28_seql alignment with homologues from *D. fasciculatum, D. purpureum* and *P. pallidum*. Upper panel, Jalview [44] multiple sequence alignment together with Jpred secondary structure prediction and its associated confidence, ‘JNETCONF’ [45]. Green arrows represent extended strands and red bars represent helical regions. In the alignment IBN_N (purple) and HEAT_EZ (red) domains are highlighted. Lower panel, MrBayes [46] phylogenetic tree annotated with SMART [47] domain architectures determined. Each amino acid in the multiple alignment is coloured according to the clustalx [48] colour scheme.

### Genomic cloning of orphan *Dictyostelium discoideum* mRNAs

The newly assembled transcripts that could not be mapped onto the genome are either contaminants or genuine mRNAs for which the genomic counterpart is in an assembly gap of the genome. To investigate the latter option, we used PCR to attempt to amplify the genes from *D. discoideum* genomic DNA (gDNA). Oligonucleotide primers were designed to amplify regions of about 0.5 – 1.4 kb of 11 transcripts (Table S3). The amplified size can however be larger due to the presence of introns. For eight transcripts, corresponding gDNAs could be amplified, but for two genes two transcripts were part of the same gene (Figure S3). The six genes in total were all protein coding genes. For three transcripts, comp470_c0_seql, comp4678_cl_seql and comp2066_cl_seql no PCR products were obtained, but the first two transcripts contained multiple stop codons in all reading frames and are likely assembly errors. The amplified PCR products were sub-cloned and sequenced from both ends. Sequences were assembled and aligned with the transcript sequence. Apart from just a few mismatches, the transcript and gDNA sequences were identical (Table S4). Only one amplified fragment contained introns (Table S4). Six out of seven of the protein coding orphan transcripts therefore had a counterpart in the genome. Overall, deciphering the genomes of organisms is a key step in being able to probe their biology. With the advent of high-throughput sequencing technologies this has become a simpler problem to solve. Yet it is still not trivial to finish a genome assembly without any gaps [37], The genome sequence on its own, however, imparts very little functional information and requires annotation of genes, transcripts and regulatory regions to be scientifically useful [7], Many gene annotation methods are dependent on either homology to related species [28, 38] or via gene finding prediction algorithms [39, 40] or ideally both. However, the first method will miss all unusual or species-specific genes, while both methods fall short of accurately predicting intron-rich genes, genes with alternative or non-canonical splice sites or genes with very short exons. The ability to generate a whole transcriptome for a given species and use it to empirically annotate the genome has the power to confirm and correct any errors introduced with other methods. This has been achieved with expressed sequence tags (ESTs) in the past [41], but now can be performed with RNA-Seq short read data [32].

This evidence-based methodology is non-trivial and is not perfect. There are examples where the data is not adequately represented in the final transcript set when interpreted by the human eye. In addition, PASA only defines protein-coding genes meaning that all non-coding RNAs (ncRNAs) will be ignored and will not be in the final annotation unless already identified in the reference. Identifying ncRNAs is difficult as they have no obvious products and well-defined sequence features [42], This does not negate their importance or relevance to the Dictyostelia.

## CONCLUSION

For the first time, In this study, we present a *de novo* transcriptome assembly in four social amoeba species and with these data we have:

- Created a final set of of 11,523 (*D. discoideum*), 12,849 (*P. pallidum*), 12,714 (*D. fasciculatum*) and 11,315 (*D. lacteum*) transcripts.
- Substantially updated the existing transcript annotations by altering models for more than half of all the annotated transcripts.
- Identified changes to thousands of transcripts in the predicted gene models of *P. pallidum*, *D. lacteum* and *D. fasciculatum* many of which affect the protein coding sequence.
- Identified and validated six novel transcripts in *D. discoideum*.
- Putatively identified dozens to hundreds of novel genes in all four species.
- Identified errors in the genome sequence of at least two *D. discoideum* genes (clcD and DDB_G0279477). With the possibility of, at least, another 104 genes having sequence errors.
- Found hundreds of putatively alternatively spliced transcripts in all species, something which has not been identified before in *P. pallidum, D. lacteum* or *D. fasciculatum*.

By combining methodologies we now have a better and more complete description of the transcriptome for these four species. This is not an end-point, however, but a further step towards fully finished genomes. More data and more manual refinement will be required to improve the annotations further.

## DECLARATIONS

### List of abbreviations

BLAT: BLAST-like alignment tool
bp: basepairs
CDS: coding sequence
CEGMA: Core Eukaryotic Genes Mapping Approach
CEG: Core Eukaryotic Genes
cDNA: coding DNA
contig: contiguous sequence
ERCC: External RNA Controls Consortium
EST: expressed sequence tag
gDNA: genomic DNA
GMAP: Genome Mapping and Alignment Program
PCR: polymerase chain reaction
mRNA: messenger RNA
ncRNA: non-coding RNA
rRNA: ribosomal RNA
RNA-seq: RNA sequencing
PASA: Program to Assemble Spliced Alignments
UTR: untranslated region

### Ethics approval and consent to participate

Not applicable.

### Consent for publication

Not applicable.

### Availability of data and material

The datasets supporting the conclusions of this article are available at figShare, http://dx.doi.org/10.6084/m9.figshare.3384364, the European Nucleotide Archive (project IDs: PRJEB12875, PRJEB12907, PRJEB12908, PRJEB12909) and as an Integrated Genome Browser [35] QuickLoad server at https://compbio.lifesci.dundee.ac.uk/Quickload/Dictyostelidassemblies.

### Competing Interests

The authors declare that they have no competing interests.

### Authors’ contributions

RS and CC developed experimental design, performed analyses and wrote the manuscript. HML performed gene amplification by PCR. PS and GJB contributed to experimental design and manuscript preparation. CS and GG prepared and sequenced RNAs, respectively.

## Funding

RS, CS, HLM and PS are funded by BBSRC grant BB/K000799/1 and Wellcome Trust grant 100293/Z/12/Z. The GSU was funded under the Wellcome Trust Strategic Award 098439/Z/12/Z.

## Acknowledgements

We thank Dr Thomas Walsh and the School of Life Sciences IT team for management of our high performance computing infrastructure and help in supporting our computation work.

